# A Chromosome-Scale Assembly of the Enormous (32 Gb) Axolotl Genome

**DOI:** 10.1101/373548

**Authors:** Jeramiah J. Smith, Nataliya Timoshevskaya, Vladimir A. Timoshevskiy, Melissa C. Keinath, Drew Hardy, S. Randal Voss

## Abstract

The axolotl (*Ambystoma mexicanum*) provides critical models for studying regeneration, evolution and development. However, its large genome (~32 gigabases) presents a formidable barrier to genetic analyses. Recent efforts have yielded genome assemblies consisting of thousands of unordered scaffolds that resolve gene structures, but do not yet permit large scale analyses of genome structure and function. We adapted an established mapping approach to leverage dense SNP typing information and for the first time assemble the axolotl genome into 14 chromosomes. Moreover, we used fluorescence *in situ* hybridization to verify the structure of these 14 scaffolds and assign each to its corresponding physical chromosome. This new assembly covers 27.3 gigabases and encompasses 94% of annotated gene models on chromosomal scaffolds. We show the assembly’s utility by resolving genome-wide orthologies between the axolotl and other vertebrates, identifying the footprints of historical introgression events that occurred during the development of axolotl genetic stocks, and precisely mapping several phenotypes including a large deletion underlying the *cardiac* mutant. This chromosome-scale assembly will greatly facilitate studies of the axolotl in biological research.

## INTRODUCTION

In the late 19^th^ century, the axolotl (*Ambystoma mexicanum*) emerged as a model for studying vertebrate development (Reiss et al. 2015) and has continued to serve as a uniquely informative model for studying diverse topics, including: regeneration (Voss et al. 2009), evolution of germline (Evans et al. 2014), genome evolution (Smith et al. 2009; Voss et al. 2011; Keinath et al. 2015; Keinath et al. 2016; Nowoshilow et al. 2018) and the origin of adaptive phenotypes (Voss and Shaffer 1997; Voss and Shaffer 2000). Recent advances in sequencing, assembly and genetic manipulation have led to a resurgence of this model and are beginning to surmount longstanding issues associated with the incredibly large size of the axolotl genome (~ 32 gigabases: Gb). The axolotl genome assembly has seen steady improvement over the last few years (Smith et al. 2005; Keinath et al. 2015; Evans et al. 2018; Nowoshilow et al. 2018). The latest version of the assembly harbors annotated models for a vast majority of axolotl genes on scaffolds that typically exceed a megabase (Mb) in length (N50 ~ 3 Mb) (Nowoshilow et al. 2018). Even with the remarkable computational methods and extensive sampling of long reads brought to bear in the completion of the most recent assembly, the typical contig represents ~1/1,000^th^ of a chromosome and contains on average 1-2 complete or fragmented genes and associated promotor/proximate regulatory sequences. Recognizing that greater contiguity will permit more in-depth analyses that leverage large-scale genomic signatures (*i.e.* identification of regulatory elements, genetic association studies and genome evolution), we sought to improve the contiguity of the genome assembly toward the goal of achieving chromosome-scale scaffolding.

Initial attempts to develop a chromosome-scale assembly using a proximity ligation approach (DoveTail) (Putnam et al. 2016) revealed limitations of the approach and yielded few useful results, despite extensive sequencing/computational costs and time associated with analyses of the resulting datasets. This led us to adapt established meiotic mapping methods for *Ambystoma* as a tool to generate dense genome-wide scaffolding information. Hybrid crosses between *A. mexicanum* and *A. tigrinum* (and other species) have been generated and used to develop meiotic maps for the species and infer the positions of quantitative trait loci (QTL), sex and Mendelian pigment mutants (Voss and Shaffer 1997; Voss and Shaffer 2000; Voss et al. 2001; Voss and Smith 2005; Smith and Voss 2006; Smith and Voss 2009; Page et al. 2013; Woodcock et al. 2017). For example, hybrid crosses have been used to dissect the genetic basis of paedomorphosis, a hallmark trait of salamanders (Voss and Shaffer 1997; Voss and Shaffer 2000; Voss and Smith 2005). Paedomorphosis reflects the loss of metamorphic changes in a derived species relative to its ancestor. The ambystomatid ancestor and several extant species, including *A. tigrinum*, undergo a metamorphosis from a larval to adult stage prior to reproduction, whereas *A. mexicanum* reaches sexual maturity in its larval form and does not undergo a metamorphosis. While other factors likely contribute to the alternate expression of paedomorphosis in *A. mexicanum* and *A. tigrinum*, hybrid mapping studies have found that much of this variation can be attributed to a single genomic region, known as the *met* QTL. In the last year, sparse meiotic mapping information was also used to identify the genes underlying two pigment mutants (Woodcock et al. 2017), however, the genes for many mutants remain unknown. Development of a chromosome-scale assembly would greatly accelerate gene discovery and make it possible to identify epigenetic changes associated with embryogenesis, post-embryonic development and regeneration.

Here we use low coverage sequence data from 48 segregants of a hybrid *A. mexicanum*/*A. tigrinum* backcross to identify and localize 12.6 million SNPs and 27.3 Gb of corresponding *A. mexicanum* sequence on 14 chromosomal scaffolds, which corresponds to the known number of chromosome pairs. Additionally, we used fluorescent in situ hybridization (FISH) mapping of BAC contigs to physically anchor scaffolds to specific chromosomes and verify that these scaffolds correspond to single (entire) chromosomes. We use this new assembly to improve resolution of orthologous regions between salamander and other vertebrates, identify regions of introgression that have persisted since the early establishment of axolotl genetic stocks, and precisely map the location of major effect loci, including sex, the *met* QTL, and a large deletion underlying *cardiac,* a classical Mendelian mutant.

## RESULTS

### SNP Typing and Linkage Analysis

Our analyses of low coverage sequence data yielded 12.6 million *A. mexicanum* / *A. tigrinum* polymorphisms segregating in the mapping panel. These were distributed across 98,802 scaffolds with individual scaffolds containing between 1 and 12,108 polymorphisms. Although sequencing depth at any position was insufficient to reliably assess genotype, we were able to generate consensus genotypes (homozygous *A. mexicanum*: or heterozygous *A. mexicanum* / *A. tigrinum*) for 69,250 scaffolds by averaging across five or more multiple adjacent SNPs. A vast majority of scaffolds were characterized by segregation ratios approximating 1:1, consistent with expectations from the backcross mapping panel and a small number with aberrant segregation ratios were flagged as potential assembly errors (Supplementary Table 1).

Consensus genotypes were used to infer the grouping and ordering of scaffolds within linkage groups. To avoid confounding effects of genotyping errors on the estimation of the recombination structure of the linkage map, we selected a subset of 8,758 scaffolds with genotypes supported by more than 200 polymorphic sites to build a framework genetic map. After manual curation, these markers yielded 14 linkage groups spanning a total of 5,364 cM (Figure 1), which is consistent with microscopic observations of the *A. mexicanum* karyotype and chiasma frequencies (Callan 1966). The average intermarker distance within this framework map was less than one centimorgan (0.6 cM), smaller than the minimum distance that can be estimated from 48 backcross offspring (~2.1 cM). Notably, the *met* QTL interval was inflated relative to the rest of the map because recombinant individuals for this region were purposely included in the mapping panel (see below). After construction of the framework map, an additional 17,819 markers were placed on the meiotic map based on similarity of genotypes to the framework map. Additional scaffolding and orientation information from previously-published RNA-Seq studies (Evans et al. 2014; Bryant et al. 2017; Caballero-Perez et al. 2018) and bulked-segregant analyses were used to tune internal ordering of scaffolds and add another 1,097 previously unplaced scaffolds (grand total of 27,674). In total, 27.5 Gb of sequence was scaffolded onto 14 linkage groups (chromosomes), with the total length of scaffolded sequence per chromosome ranging from 3.14 to 0.66 Gb (Figure 1, Supplementary Tables 2 and 3). By comparison, the entire human genome is approximately 3.2 Gb.

**Figure 1.**
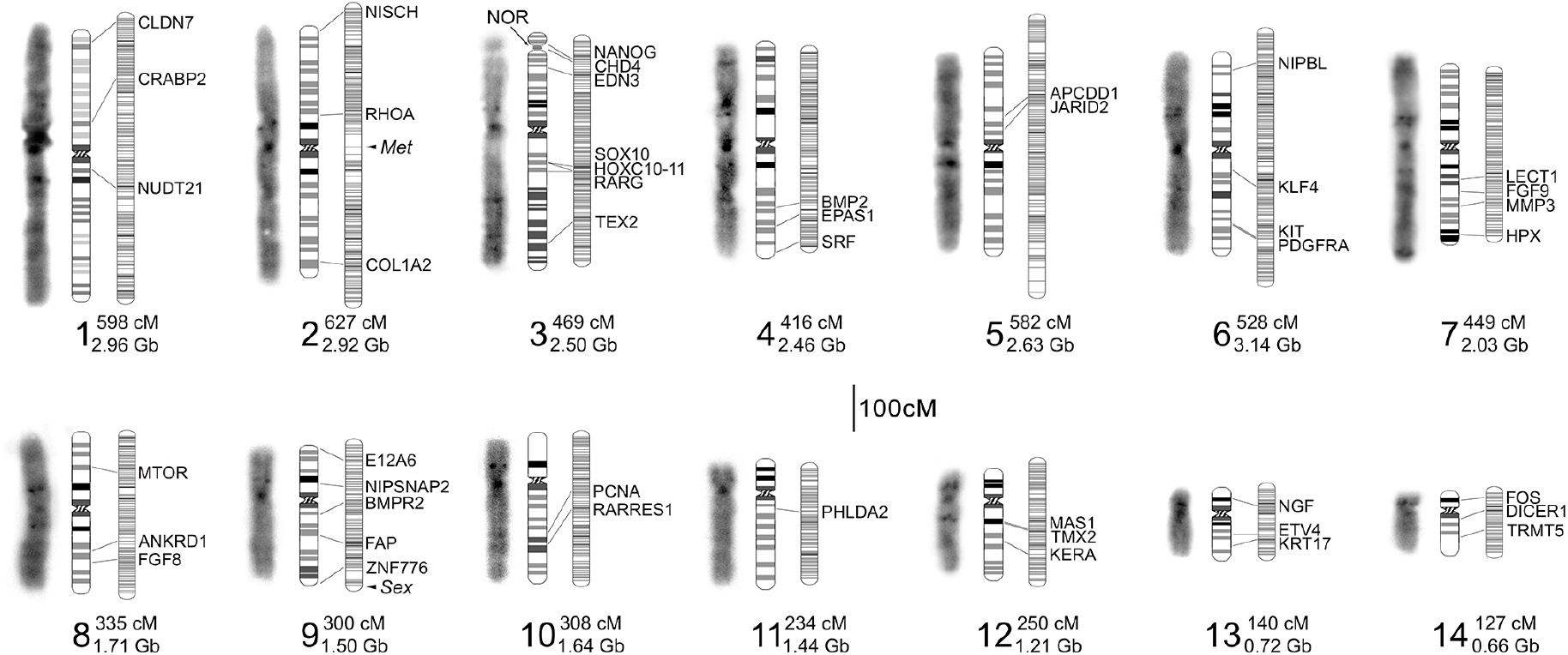
Summary of assembled chromosomes from the *A. mexicanum* genome. Three images are shown for each chromosome (1 – 14): left - microscopic image illustrating DAPI-banding, center - an ideogram summarizing banding patterns and relative sizes of chromosomes from several spreads (see also Supplementary Figure 1), right – the corresponding linkage group. Groups of linked markers at a given position are represented by horizontal marks on linkage groups. The location of genes and hybridization signals from gene-anchored BACs are labeled with the corresponding gene ID and connecting lines. The length of each linkage group and assembled chromosome is given next to its numerical label. The locations of three additional features are highlighted: *met* – a major quantitative trait locus controlling timing and incidence of metamorphosis, *sex* – the sex determining locus, and NOR – the nucleolar organizer, which harbors most ribosomal RNA copies.

To validate the large-scale structure of our scaffolded genome and more directly assign linkage groups to chromosomes, we performed FISH analyses using a collection of 46 gene-anchored BACs. These hybridizations confirmed that our linkage groups correspond to individual chromosomes, confirmed the large-scale ordering of scaffolds along the length of chromosomes, and allowed us to assign assembled scaffolds to karyotypically definable entities (Figure 1, Supplementary Figure 1).

### Conservation of synteny and chromosome evolution

To further assess the degree to which our scaffolded genome recovers conserved gene orders across vertebrates, we aligned predicted *A. mexicanum* proteins to those annotated in the genomes of chicken, human and frog (*Xenopus tropicalis*: XT). These revealed broad scale conservation of genome structure, and fusion/fission events that underlie differences in chromosome number and content between taxa (Figure 2, Supplementary Tables 4 - 6). In general, these confirm patterns reported from earlier mapping studies, revealing that *X. tropicalis* and *A. mexicanum* (AM) genomes are composed of ancestral chromosomes and microchromosomes that were independently fused after the separation of frog and salamander lineages. These also confirm that conserved regions corresponding to chicken chromosomes (GG) Z and 4 were linked in the ancestral amphibian (Smith and Voss 2006;Voss et al. 2011;Keinath et al. 2016) and identify two other conserved linkages (portions of GG8/11 homologous to AM1/XT4, and portions of GG6/21 homologous to AM8/XT7), which represent previously unreported fusions events that likely occurred in the common ancestral lineage of all extant amphibians. Moreover, the extensive conservation of synteny revealed by these analyses serves as additional support for the accuracy of the large-scale structure of the assembled chromosomes.

**Figure 2.**
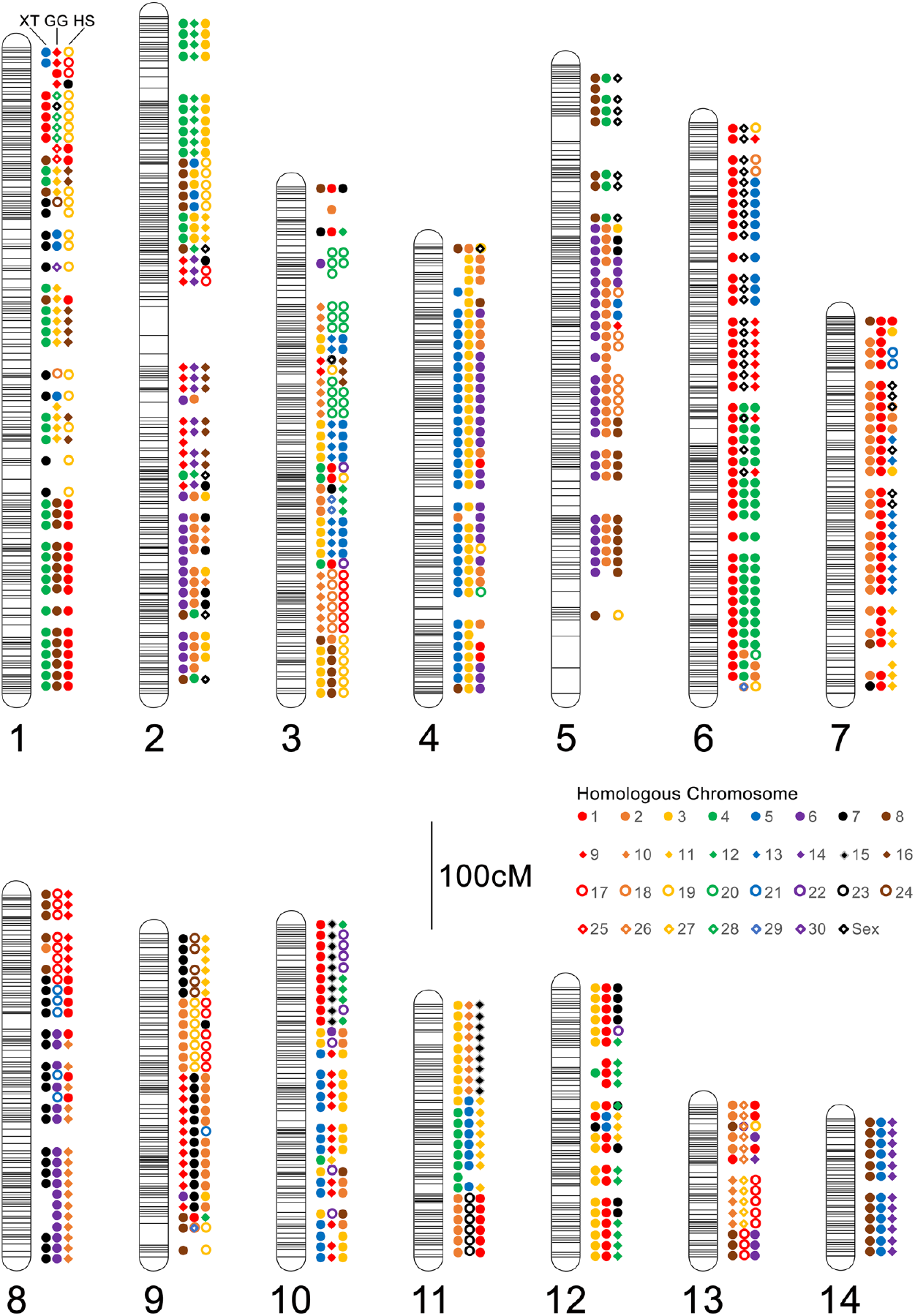
Patterns of conserved Synteny across assembled *A. mexicanum* chromosomes. Colored shapes represent the consensus homologous chromosomes (see inset legend) for each 10 cM interval. For each chromosome, homologs are shown in a consistent order from left to right: left - *X. tropicalis* (XT), center - chicken (*Gallus gallus*: GG), right - human (*Homo sapiens*: HS). *A. mexicanum* chromosome are designated by numerical labels corresponding to those in Figure 1.

### Distribution of polymorphisms

A recent study showed that the vast majority of *A. mexicanum* used in biological studies derive from an *A. mexicanum / A. tigrinum mavortium* cross that was performed in 1962 (Humphrey 1967). This cross was performed to introgress an albinism gene (*tyr*) into a captive population to create a new mutant strain. Analysis of SNPs across the axolotl genome reveal that the density of polymorphisms (in 1 Mb windows) is relatively consistent, averaging 0.43 polymorphisms per kilobase (s.d. = 0.20, N = 27,518 regions). However, the fingerprint of the *tyr* introgression is evident when examining density distributions for typed polymorphisms (Figure 3, chromosome 7). The density of polymorphisms is lower at the position of *tyr* because more tigrinum-like sequence has remained in linkage disequilibrium within this region. Notably, other regions show similar depression in SNP densities, including one region near the centromere of the sex chromosome and another large region on chromosome 1. Notably, the kisspeptin-54 receptor (KISS1R) is located near the center of the presumptive *A. tigrinum* introgression on chromosome 1. This gene is a key regulator of gonadotropin-releasing hormone signaling and variation in this gene is known to be associated with the onset timing of sexual maturity in mammals (Seminara et al. 2003;Popa et al. 2008). We speculate that this and other retained *A. tigrinum* mavortium DNA sequences were retained because they confer fitness benefits in the context of laboratory culture and husbandry practices.

**Figure 3.**
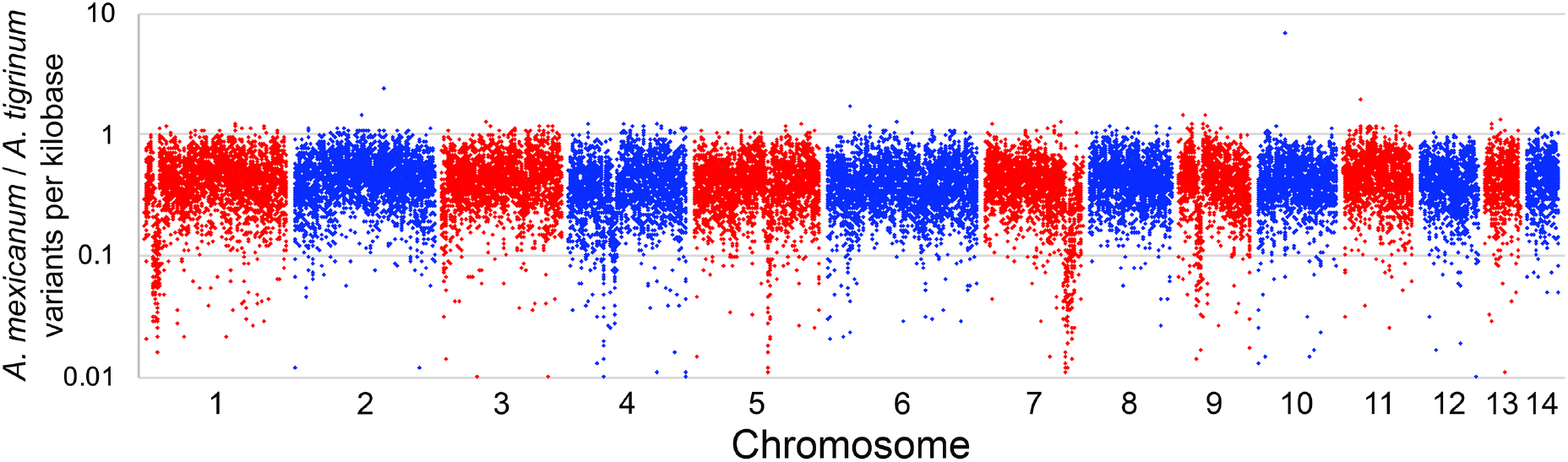
Plot showing the distribution of segregating variants within the mapping panel used to generate scaffolding information. The density of polymorphisms within 1 Mb intervals is relatively uniform, however, a few intervals show substantially reduced densities, including regions on chromosomes 1, 5, 7 and 9.

### The Genetic Basis of the cardiac Mutant

To further illustrate the utility of the assembly in performing genetic analyses in laboratory stocks of *A. mexicanum*, we sought to localize the mutation underlying the classical Mendelian mutant *cardiac* (*c*) (Humphrey 1972). Due to the large size of the genome and the cost associated with generating sufficient data to accurately assess genotypes from DNA sequencing data, we drew from successful efforts in newt (*Notophthalmus viridescens*) and used transcriptomic data to efficiently sample gene-anchored polymorphisms (Keinath et al. 2016). This sampling strategy effectively reduces the size of the genome and more deeply subsamples the ~100 Mb of largely, gene-anchored sequences that are processed into mature transcripts. To further reduce costs associated with localizing the mutant, we used a bulked-segregant mapping strategy, sequencing RNAs from two pools of embryos that exhibited the *c* phenotype and two pools of their wildtype siblings (N = 10 per pool). Genotype/phenotype associations calculated from these data revealed a strong peak near the predicted location of the centromere of chromosome 13 (Figure 3). This is consistent with early gynogenetic diploid mapping experiments that predicted tight linkage between *c* and it’s centromere, but did not identify the specific chromosome carrying the *c* mutation (Armstrong 1984). Manual inspection of the region under this peak revealed that troponin 2 (*tnnt2*), a cardiac-specific troponin, was located under the association peak on chromosome 13, though notably *tnnt2* was not sampled at sufficient depth in our RNA-Seq data to permit calculation of test statistics for the gene itself. To further assess this association, we performed genomic PCR using primers targeted to *tnnt2* exons. These revealed that all exons could be amplified from wildtype individuals, as well as a specific and reproducible lack of amplification of exons 8 and 9 from *cardiac* homozygotes (Figure 4c). We consider this strong evidence that this specific deletion within *tnnt2* underlies the *c* mutation.

**Figure 4.**
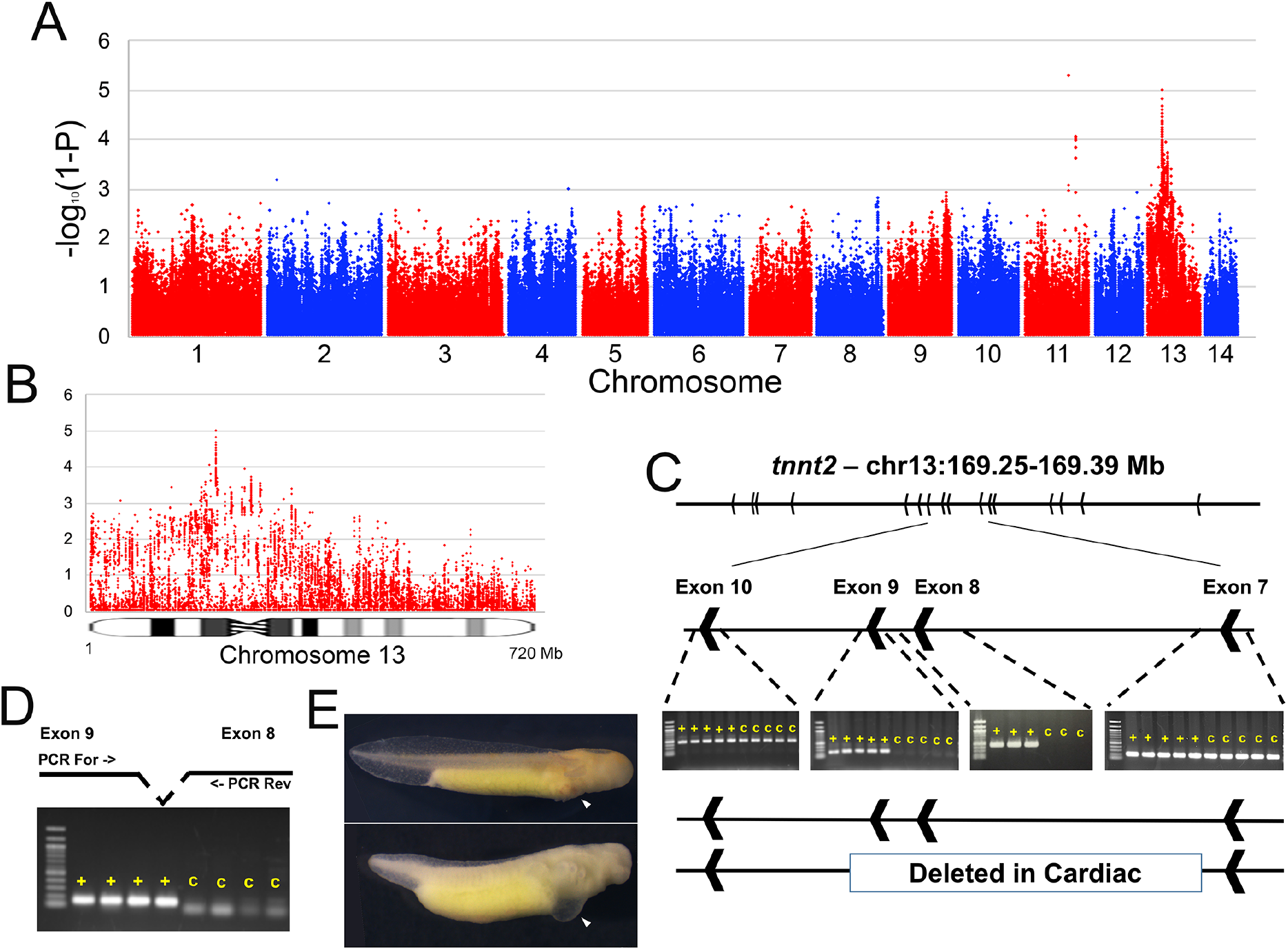
Localization and characterization of the *cardiac* mutation. A) The genome-wide distribution of P-values for tests of association between transcribed polymorphisms and the *c* mutant phenotype. B) Distribution of P-values across chromosome 13. C) Identification and PCR validation of a large deletion that is associated with the *c* phenotype and removes exons 8 and 9 of *tnnt2*. D) Verification that these same exons are expressed in wildtype individuals, but not in individuals with the *c* phenotype. E) Example of two siblings from a cross segregating for the *cardiac* mutation; top – wildtype, bottom cardiac homozygote. Note the thoracic edema and lack of erythrocytes in the developing heart (white arrows) of cardiac mutants.

## DISCUSSION

This study presents a relatively cost-effective approach to scaffolding the large axolotl genome and validates the resulting assembly using both FISH and analysis of synteny conservation across vertebrates. We also demonstrate the utility of this assembly in resolving patterns of genome-wide SNP variation that 1) reveal the footprints of a historical introgression event and 2) resolve the genetic underpinnings of the classical mutant phenotype *cardiac* using a simple RNA-Seq based approach that efficiently navigates the large axolotl genome. These analyses set the stage for routine analyses at this scale, including not only SNP-based assays, but other analyses that involve large-scale signatures such as epigenetic remodeling events and promotor-enhancer interactions that will be critical for dissecting the regulatory logic of *A. mexicanum* development and regeneration.

### TNNT2 and *cardiac*

The *cardiac* mutation was first described 45 years ago (Humphrey 1972) and has been the subject of numerous descriptive, biochemical and genetic analyses. Animals carrying homozygous recessive alleles for *c* lack organized cardiac myofibrils, do not generate a heartbeat and die around the time of hatching (Humphrey 1972;Lemanski 1973). This mutation has been attributed to defects in the expression of tropomyosin (Zajdel et al. 1999), troponin (Sferrazza et al. 2007;Zhang et al. 2007) and non-coding RNA (Lemanski et al. 1996;Zhang et al. 2003;Kochegarov et al. 2013), and has been associated with a point mutation in the noncoding myofibril-inducing RNA (MIR) (Zhang et al. 2003). Our analyses provide strong evidence that the genetic basis of the mutation lies in a large deletion that resulted in the loss of internal *tnnt2* exons. Notably, the interval does not contain the myofibril-inducing RNA, which lies on chromosome 12 upstream of the Cholinergic Receptor Muscarinic 2 (CHRM2) gene. It seems plausible, however, that cis modulation of cardiac contractile activity by CHRM2 might underlie improvement in cardiac function observed in experiments dissecting the effects of myofibril-inducing RNA.

The verification of linkage between *cardiac* and the centromere highlights the potential of bulked-segregant analyses in axolotl and earlier approaches that were used to map genes prior to the advent of modern molecular methods. The availability of a chromosome-scale assembly will dramatically facilitate forward genetic screens of additional mutants and allow for more accurate and precise design of gene-editing tools to empower reverse genetic experiments.

### Localization of the Major Effect QTL: met

The mapping cross used for this study was originally designed to dissect the genetic basis of paedomorphosis in *A. mexicanum*, whereby this species reaches sexual maturity without undergoing metamorphic changes that are typical of most amphibian species. Many of the individuals selected for this study were known to possess recombinant genotypes near the *met* locus. As such, estimated recombination frequencies were inflated near the position of the *met* QTL (Figure 1, chromosome 2). This inflation permitted us to more precisely define the limits of *met* and identify a relatively small (~10 Mb) genomic region containing 9 annotated genes, including homologs of NGFR, SETD2, KIF9, KLHL18 and 5 predicted loci (LOC101951429, LOC109139901, LOC102363594, LOC106731944, LOC102943813). It seems possible that variation in one or more of these genes may regulate the expression of metamorphic vs. paedomorphic modes of development, or alternately, that a gene (present in other metamorphic species) may have been lost within this region of the *A. mexicanum* genome. Future efforts to sequence and assemble additional salamander genomes should shed light on the ancestral structure of this region and changes that are associated with life history evolution.

### Broad Utility in Axolotl and Other Salamanders

We anticipate that the improved assembly presented here will facilitate diverse studies in urodeles by providing insights into genome structure that were not previously accessible in any salamander species. Long-range interactions are critical for regulating transcription, particularly in large genomes and even in the comparatively small human genome. Enhancer/promotor interactions may involve looing structures that arch over multiple intervening genes and span megabases (Whalen et al. 2016;Cao et al. 2017). Resolving such interactions will likely be critical to understanding the gene-regulatory basis of tissue regeneration. This assembly should also be useful for interpreting patterns from large-scale sampling of polymorphisms from species and natural populations. Thus far, it has been difficult to effectively control for linkage when estimating demographic parameters and genome-wide patterns of divergence (McCartney-Melstad et al. 2016;Rodriguez et al. 2017;Nunziata and Weisrock 2018), particularly given evidence for relatively strong conservation in genome structure between *Ambystoma* and newt (Keinath et al. 2016), which diverged ~150 million years ago. Extension of similar approaches to species without existing methods for production of hybrid crosses should also be relatively straight-forward, provided offspring or other meiotic products can be sampled from individuals with modest levels of heterozygosity.

## METHODS

### Sequencing of Segregants for Meiotic Mapping

DNA was extracted from 48 individuals that were selected from a previously described mapping population that was generated by backcrossing F1 hybrid *A. mexicanum*/*A. tigrinum* to *A. mexicanum* (Voss and Smith 2005). Individuals were selected to include individuals of known sex and that were previously inferred to have inherited recombinant genotypes near the *met* locus (Voss and Smith 2005). DNA was extracted via phenol-chloroform extraction (Sambrook and Russell 2001). Outsourced library prep and Illumina sequencing (HiSeq2500 V4, 125bp paired-end reads) was performed by Hudson Alpha Genome Services Laboratory. In total these yielded ~14 billion reads, with each individual being sequenced to ~1X coverage.

### SNP Typing

Sequence data were aligned to the genome assembly (ambMex3) using BWA-mem (Li and Durbin 2009), filtered by samtools view (with option -F2308) (Li et al. 2009 and processed by GATK V3.5 (McKenna et al. 2010; Van der Auwera et al. 2013) to identify variant sites and tabulate the number of reads supporting reference vs alternate alleles in each individual. Sites were then filtered to extract those that were homozygous for the reference base in a re-sequenced *A. mexicanum* (Ambystoma Genetic Stock Center ID# 130001.3, sequenced to 20X depth of coverage) and homozygous for the alternate base in a re-sequenced *A. tigrinum* (sequenced to 3X depth of coverage) in order to identify a subset of sites that are likely to differ consistently between the species. Prior to inferring genotypes, polymorphisms were filtered using VCF tools (Danecek et al. 2011) to retain a subset of individual biallelic SNPs that were spaced at a minimum distance of 75bp, characterized by observed *A. tigrinum* allele frequencies within the range expected for the backcross design (0.25 ± 0.15), and sampled in 18 or more individuals. Low-coverage SNP calls were accumulated across the length of each scaffold in order to infer whether each individual was homozygous *A. mexicanum* or heterozygous *A. mexicanum*/*A. tigrinum* by summing the number of reads that could be assigned to *A. mexicanum* or *A. tigrinum* and assessing the resulting ratios (*A. tigrinum* reads / *A. tigrinum* + *A. mexicanum* reads) to define a consensus genotype for the scaffold. Individuals with ratios reflecting no or low representation of *A. tigrinum* alleles (<0.20) were scored as homozygous *A. mexicanum*, and individuals with ratios approximating 50% *A. tigrinum* alleles (0.25 - 0.75) were scored as heterozygous *A. mexicanum*/*A. tigrinum*. Individuals with intermediate frequencies or low coverage (fewer than 4 reads) were scored as missing genotypes.

### Linkage Analysis

A meiotic map was generated from a set of 26,577 genotyped scaffolds using JoinMap4.1 (Stam 1993). A high-confidence backbone map was generated using a subset of 8,758 scaffolds with genotypes supported by more than 200 polymorphic sites. This map was manually curated by examining all intervals greater than 10 cM, recursively removing loci, and examining the effect of removal on map distance. Markers that increased map distance were considered erroneous genotypes and were removed from the backbone map. This backbone map was enriched with additional scaffolds with genotypes that were identical or highly similar (Jaccard distance < 10%) to backbone scaffolds, without impacting the relative ordering or estimated recombinational distance between high-confidence markers on the backbone map.

### Ordering and Orientation with RNAseq data

Paired-end RNA-Seq datasets [from NCBI BioProjects PRJNA300706; PRJNA306100; PRJNA312389; PRJNA354434] and bulked-segregant mapping studies (below) were aligned to genomic scaffolds and predicted gene models from ambMex3 using BWAMEM (Li and Durbin 2009), and the resulting alignments were processed by Rascaf (Song et al. 2016) and AGOUTI (Zhang et al. 2016) to identify linkages that were supported by ≥5 read pairs (minimum mapping quality 20 with no mismatches for AGOUTI). The resulting linkage and orientation information was appended to the linkage map using ALLMAPS (Tang et al. 2015).

### Karyotypic Analyses and FISH Mapping

Suspensions of mitotic cells for chromosome preparations were made according to previously described protocol (Keinath et al. 2015). Cells were applied to a cold glass slide and immediately placed in humid chamber at 60°C in order to spread chromosomes, air-dried, aged overnight at 65°C and held at −20°C in 100% ethanol until processing. To develop idiograms, DAPI stained early metaphase chromosomes were imaged, grayscaled, inverted and contrasted using Adobe Photoshop (Demin et al. 2011). Chromosome images were straightened using ImageJ (https://imagej.nih.gov/ij/) and aligned for comparison. Two gradations of gray were used to depict reverse DAPI banding patterns: strong DAPI-positive puncta (bands that do not necessarily span the breadth of large chromosomes) (Rudak and Callan 1976) are designated by black bars, other stronger and fainter bands are respectively designated by dark and light gray bands in ideograms (Supplementary Figure 1).

For hybridization of individual genes, BAC libraries were screened by PCR as previously described (Smith et al. 2009) in order to identify a set of 46 BACs that correspond to known *A. mexicanum* genes (Supplementary Table 7). DNA from individual BAC clones was isolated using the Qiagen Large Construct kit (Qiagen Science, Germantown, MD, USA); or via an outsourced purification service provided by Clemson University Genomics Institute. FISH was performed as previously described for BAC-DNA probes with Cot_2_DNA suppression (Timoshevskiy et al. 2012). Two color fluorescence *in situ* hybridization was used for localization signals from BAC-related markers on axolotl chromosomes. After localization of each new marker to a specific chromosome, according to size and DAPI banding patterns, markers on each particular chromosome were cohybridized to define their relative location and ordering.

### Comparative Synteny Analyses

Annotated genes were aligned (blastp) (Altschul et al. 1990; Camacho et al. 2009) to Human (*Homo sapiens* Refseq proteins from hg38), Chicken (*Gallus gallus* Ensembl proteins from galGal4) and *Xenopus tropicalis* (v9.0) proteins. All homologs with a bitscore >99% of the best match and ≥100 were considered homologous and integrated into comparative maps. Orthologous segments were defined across the map by tabulating the corresponding chromosomal location of genes encoding presumptively orthologous proteins and calculating the consensus across all 10 cM bins.

### Sequencing of Bulked Segregants

#### Preparation and sequencing of RNA pools

A cross [[Ambystoma Genetic Stock Center (AGSC - RRID:SCR_006372) spawn ID: 14427) was generated between two heterozygous *cardiac* carriers [AGSC IDs: 13911.2 (male) and 13970.1 (female)] and embryos were raised in 10% Holtfretter’s solution (Armstrong et al. 1989) at 20°C until completion of early cardiac development (for wildtype embryos - RRID:AGSC_100E). A total of 20 *cardiac* mutants (RRID:AGSC_104E) and 20 wildtype embryos were sampled for RNA extraction. Individual embryos were dissociated with 23- and 26-gauge needles and RNA was isolated with Trizol and then further purified using a Qiagen RNeasy Mini Kit with DNAse treatment. RNA extracts were pooled in groups of 10 (two pools of wildtype and two pools of *cardiac* homozygotes) in equimolar ratios. These RNAs were used to generate outsourced Poly A RNA-Seq libraries and were sequenced on an Illumina HiSeq 2500 (V4 sequencing chemistry, 125bp paired-end reads) by Hudson Alpha Genome Services Laboratory.

#### Analysis of RNAseq datasets

Reads from each pool of *cardiac* homozygotes or wildtype individuals were mapped to the complete set of annotated genomic gene models (Nowoshilow et al. 2018) using BWA-MEM (Li and Durbin 2009). Alignments were post-processed to remove duplicate reads using picard-tools-1.97 (http://broadinstitute.github.io/picard) and raw SNPs were identified for the complete set of transcriptome sequences with a putative human or chicken homolog and all genomic gene models using GATK V3.5 (McKenna et al. 2010;Van der Auwera et al. 2013). SNPs were filtered to identify sites that segregated two alleles in all four pools within each cross, and that were supported by 20 or more independent reads.

#### Statistical assessment of association

Association between mutant phenotypes and segregating SNPs was expressed as the absolute difference in the frequencies of the non-reference allele in pools of *cardiac* homozygotes vs wildtype individuals (Δ). SNPs that are not associated with a mutation should segregate randomly, whereas SNPs that are tightly linked to a mutation should be homozygous in affected individuals (*cardiac* homozygotes) and heterozygous in 2/3 of unaffected (wildtype) individuals. This value has a theoretical range between zero and one, with an expected value of 0.6̅ for a perfectly associated SNP. Probability values for association statistics were calculated based on the normalized (√ transformed) distribution of observed Δ values (Supplementary Figure 2).

## DATA ACESS

Sequence data for this study are published under SRA accession: SRP151479 (reviewer metadata link: ftp://ftp-trace.ncbi.nlm.nih.gov/sra/review/SRP151479_20180719_114414_cb5ae17636e975f9bf71ddf5bc542075); Genome Assembly: PRJNA482115; and as an assembly hub accessible through the UCSC genome browser (http://genome.ucsc.edu/cgi-bin/hgTracks?genome=Amex_PQ.v4&hubUrl=http://salamander.uky.edu/hubExamples/hubAssembly/Amex_PQ.v4.HUB/hub.txt).

## ACKNOWLEDGMENTS

This work was funded by grants from the National Institutes of Health (NIH) (R24OD010435) and Department of Defence (DOD) (W911NF1110475) to SRV. Animals used in this study were provided by the Ambystoma Genetic Stock Center, which is currently funded by the NIH (P40OD019794) and previously by the National Science Foundation (NSF) (DBI-0951484) to SRV. The contents of this paper are solely the responsibility of the authors and do not necessarily represent the official views of NIH, DOD or NSF. Partial computational support was provided by The University of Kentucky High Performance Computing complex.

## AUTHOR CONTRIBUTIONS

JJS and SRV conceived of the study. NT and JJS, VAT, SRV, DH, MCK contributed analyses. JJS and SRV authored the manuscript with the assistance of NT and VAT. DH and MCK provided additional edits.

